# How to predict binding specificity and ligands for new MHC-II alleles with MixMHC2pred

**DOI:** 10.1101/2023.12.18.572125

**Authors:** Julien Racle, David Gfeller

## Abstract

MHC-II molecules are key mediators of antigen presentation in vertebrate species and bind to their ligands with high specificity. The very high polymorphism of MHC-II genes within species and the fast-evolving nature of these genes across species has resulted in tens of thousands of different alleles, with hundreds of new alleles being discovered yearly through large sequencing projects in different species. Here we describe how to use MixMHC2pred to predict the binding specificity of any MHC-II allele directly from its amino acid sequence. We then show how both MHC-II ligands and CD4^+^ T-cell epitopes can be predicted in different species with our approach. MixMHC2pred is available at http://mixmhc2pred.gfellerlab.org/.

## 1. Introduction

Class II major histocompatibility complex (MHC-II) molecules play an important role in adaptive immune surveillance. MHC-II genes are highly polymorphic (***1, 2***) and have undergone fast evolution (***3–5***), leading to tens of thousands of alleles among vertebrate species. In human, there are three main gene loci (referred to as HLA-DR, HLA-DP and HLA-DQ) coding for MHC-II, all located on chromosome 6. MHC-II are expressed mostly in antigen presenting cells (e.g., B cells, dendritic cells, macrophages) but their expression can be triggered by IFNγ in other cells.

MHC-II form heterodimers (hereafter referred to as “MHC-II alleles”) composed of an alpha chain (e.g., HLA-DPA1) and a beta chain (e.g., HLA-DPB1). MHC-II alleles present short peptides on the cell surface (Figure 1A). These peptides can be specifically recognized by CD4^+^ T cells. MHC-II ligands bind to MHC-II molecules through a 9-mer binding core sequence, and flanking residues can extend outside of it on both sides (Figure 1A). Ligands are usually between 12 and 21 amino acids long with a peak in the length distribution around 15 amino acids. In general, the binding specificity of a given MHC-II allele can be accurately described by a motif (Figure 1A). In some cases, two motifs have been used, especially for alleles accommodating ligands binding in two different orientations (***6, 7***) (Figure 1B). Mathematically, MHC-II binding motifs are encoded as Position Weight Matrices (PWMs). As a result of the high polymorphism of most MHC-II genes, binding motifs show high diversity across MHC-II alleles (Figure 1C). Understanding the specificity of MHC-II molecules is key to predict potential epitopes that can be recognized by CD4^+^ T cells in viral/bacterial infection, cancer, autoimmunity and allergy.

**Figure 1:**
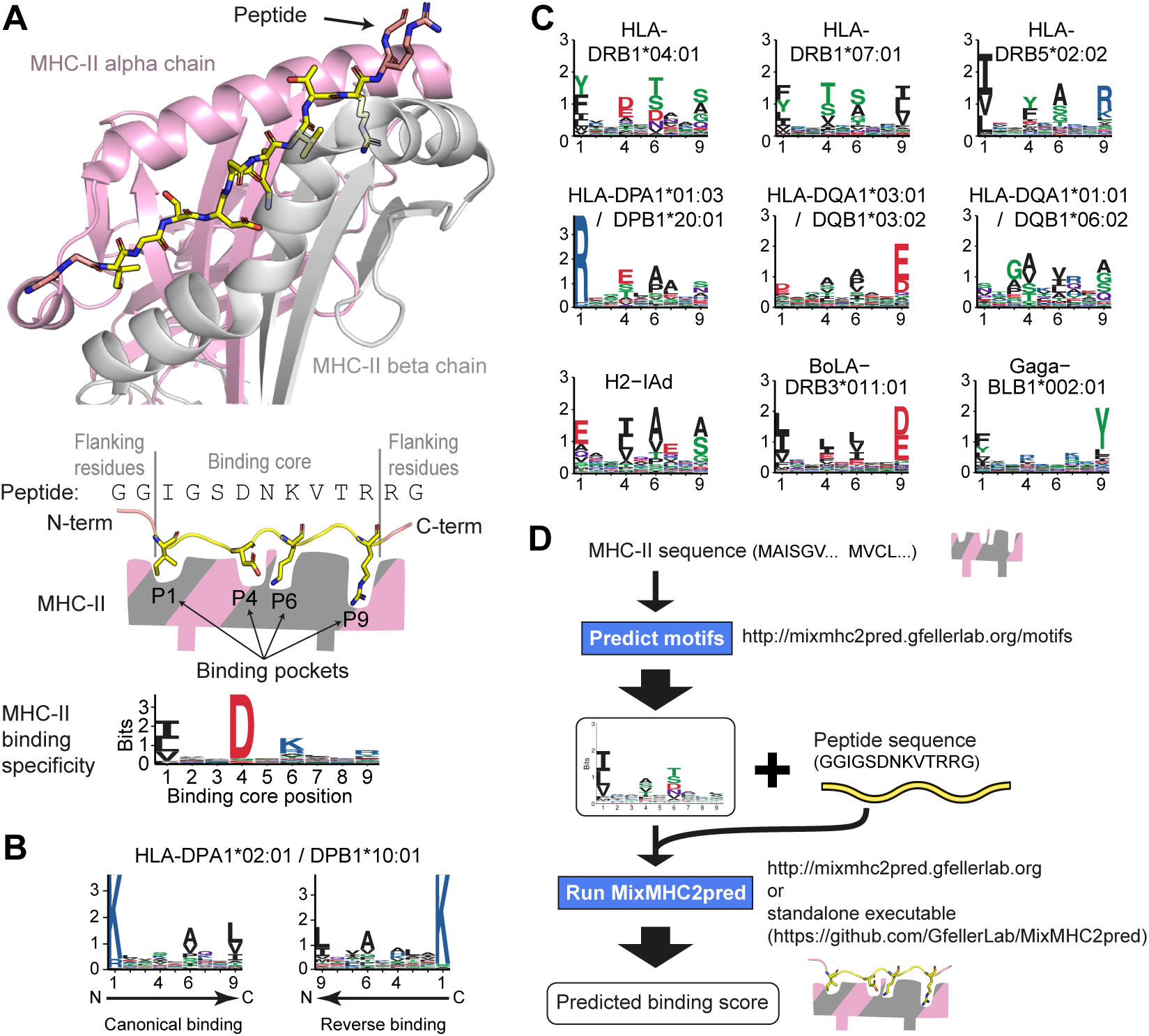
MHC-II molecules present diverse sets of short peptides. **(A)** Representative crystal structure of an MHC-II dimer (HLA-DRA*01:01/DRB1*03:01) in complex with a peptide (derived from PDB ID 7N19 (**27**)). The binding core of the peptide is shown in yellow, the peptide flanking residues in salmon, the MHC-II alpha chain in pink and the MHC-II beta chain in gray. Anchor positions P1, P4, P6, and P9 point towards the different MHC-II binding pockets and are visible in the binding motif below. **(B)** Binding specificity for HLA-DPA1*02:01/DPB1*10:01 described by two motifs corresponding to ligands binding in the canonical or reverse orientation. The arrow depicts the ligand binding orientation from P1 to P9 binding pockets, highlighting that the P1 pocket is for example on the C-terminal side of reverse binding ligands. **(C)** Example of binding motifs from various MHC-II alleles (human: HLA-xx; mouse: H2-xx; cattle: BoLA-xx; chicken: Gaga-xx). **(D)** Workflow for predicting binding specificities and ligands with MixMHC2pred. Starting from the amino acid sequence of an MHC-II allele (both alpha and beta chain sequences), its motif is first predicted (section 3.1). This motif can then be combined with sequences of putative ligands (i.e., peptides) to predict their presentation by the MHC-II allele given in input (section 3.2).

**Figure 2:**
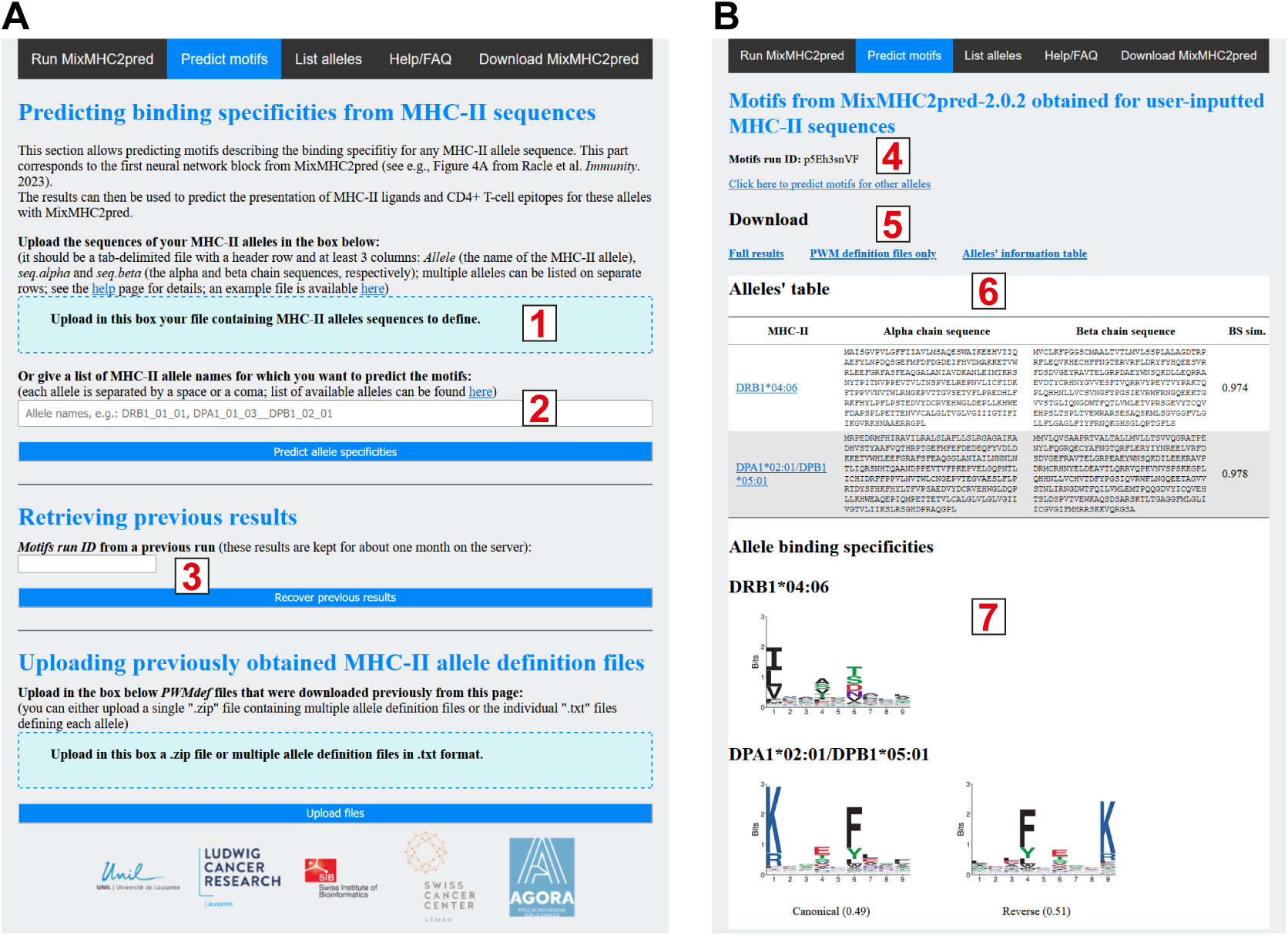
Predicting motifs with MixMHC2pred webserver. **(A)** Input page obtained from the “*Predict motifs*” of the top navigation bar allowing users to input a file with MHC-II sequences (1), input an MHC-II allele name (2) or retrieve previous results (3). **(B)** Results page, indicating the Motifs run ID (4), allowing users to download the results (5), providing information on the MHC-II sequences that have been used for motif predictions (6) and showing the predicted motifs (7).

Recent developments in MHC-II peptidomics have made it possible to obtain thousands of MHC-II ligands per sample in a single experiment in a high-throughput manner (***6–18***). Together with the development of powerful motif deconvolution algorithms to assign peptides to their cognate MHC-II alleles and identify peptide binding cores (***12, 19***), these data have enabled us and others to characterize the binding specificity of close to 100 alleles (***6, 18, 20, 21***). As of today, most of the common MHC-II alleles in human have well defined experimental motifs, but the coverage of experimental binding specificities falls short of the full diversity of human alleles (>10’000 alleles, as of November 2023 in IPD-IMGT/HLA database (***1***)). Moreover, the vast majority of MHC-II peptidomics studies were performed with human cells and much fewer MHC-II peptidomics data exist for other species. For instance, in our recent compilation of MHC-II peptidomics data, we could obtain motifs only for 4 mouse, 7 cattle and 2 chicken alleles, and nothing for other species (***6***).

This limited coverage of the MHC-II alleles in terms of experimentally determined binding motifs both in human and other species has motivated the development of tools that can make predictions of MHC-II ligands even for alleles without any known ligand (so called pan-allele predictions) (***6, 21***). These tools capitalize on existing data to learn patterns in MHC-II sequences that can predict the preference for specific ligands. In our recent work, we have shown that our pan-allele predictor MixMHC2pred was able to confidently predict the binding specificity and epitopes for any human MHC-II allele, even when the allele was absent from the peptidomics data. We have further shown that MixMHC2pred was able to perform accurate predictions of MHC-II ligands in non-human species (***6***). However, automated tools with publicly available interfaces to directly predict MHC-II binding motifs for any MHC-II allele are still missing.

In this chapter, we describe a new interface of MixMHC2pred that can be used to obtain MHC-II motifs for any MHC-II allele, starting from its alpha and beta chain amino acid sequences (Figure 1D). This chapter also explains how ligands and epitopes can be predicted for the corresponding MHC-II alleles. In section 2, we describe the *Materials* that are needed for such predictions (i.e., where and how to obtain the tools needed for the analyses). Section 3 describes step-by-step the procedure to follow: in section 3.1 to determine the binding motif of any MHC-II allele, and in section 3.2 to determine ligands and CD4^+^ T-cell epitopes for these alleles. This section is complemented by example results from analyses that could be performed. Section 4 lists *Notes* referred from the Section 3, giving some more details on specific aspects of the procedure. Finally, we conclude with a discussion in section 5, recapitulating the observations and discussing the advantages and limitations of such predictions.

## 2. Materials

Most of the results discussed in this work are based on MixMHCpred-2.0 (***6***). MixMHC2pred is available as a webserver (http://mixmhc2pred.gfellerlab.org). The part of MixMHC2pred to predict MHC-II ligands and CD4^+^ T-cell epitopes is also available as an executable with pre-compiled versions for Windows, Mac OS and Linux (https://github.com/GfellerLab/MixMHC2pred). We will outline the use of these two interfaces.

### 2.1. Hardware

There is no limitation on the hardware or operating system as MixMHC2pred can be run through a web application.

### 2.2. Software

For the main analyses and predictions, no software other than MixMHC2pred is needed. It is available as a standalone executable (precompiled for most operating systems) and webserver (tested to work with the most common browsers). For a comparison of the predicted binding specificities against motifs obtained directly from experimental data, we will also use *MoDec* whose installation is described below.

#### 2.2.1 MoDec

*MoDec* stands for <u>Mo</u>tif <u>Dec</u>onvolution (***12***) and it was developed to find motifs describing the binding specificities of the MHC-II alleles present in MHC-II peptidomics samples. It does not require any prior knowledge about potential binding specificities, and it manages to find highly reproducible motifs present anywhere along the peptide sequences, working also well in multi-allelic samples where multiple motifs are to be found.

The installation of MoDec tool is optional, since it is only needed for the supplemental comparisons showed here or to analyze some other MHC-II peptidomics data. It can be obtained from https://github.com/GfellerLab/MoDec. Installation instructions can be found in the accompanying README document.

### 2.3. Getting MixMHC2pred

#### 2.3.1 Web application

To run MixMHC2pred web application, go to the url: http://mixmhc2pred.gfellerlab.org.

#### 2.3.2 Executable

The latest release of MixMHC2pred can be obtained from https://github.com/GfellerLab/MixMHC2pred/releases/latest. Download the corresponding *zip* file (*MixMHC2pred-2.0.zip*) and uncompress it in the directory of your choice. You can then open a terminal window (or command prompt) and go inside this directory. Finally, you can verify that MixMHC2pred works by typing the following command (if on Windows):

~~~
MixMHC2pred.exe -i test/testData.txt -o test/out.txt -a
  DRB1_15_01 DRB5_01_01 DPA1_02_01 DPB1_01_01
  DQA1_01_02 DQB1_05_01 DQA1_01_02 DQB1_06_02
~~~

On Mac OS, replace *”MixMHC2pred.exe”* by *”./MixMHC2pred”* in above’s command. On Unix, replace *“MixMHC2pred.exe”* by *”./MixMHC2pred_unix”*.

The file *test/out.txt* should be the same than *test/out_compare.txt*. More detailed installation instructions can be found in the *“README.pdf”* file located in MixMHC2pred’s directory.

### 2.4. MHC-II sequences

Many MHC-II amino acid sequences across multiple species are available at the IPD-MHC database (***2***) (https://www.ebi.ac.uk/ipd/mhc). For human, MHC-II sequences are available at the IPD-IMGT/HLA database (***1***) (https://www.ebi.ac.uk/ipd/imgt/hla). To exemplify various use-cases of MixMHC2pred, we collected a few MHC-II sequences from human and non-human species from these databases (Tables 1 and S1). We will demonstrate how to predict MHC-II motifs as well as MHC-II ligands and CD4^+^ T-cell epitopes for the corresponding MHC-II alleles.

**Table 1:**
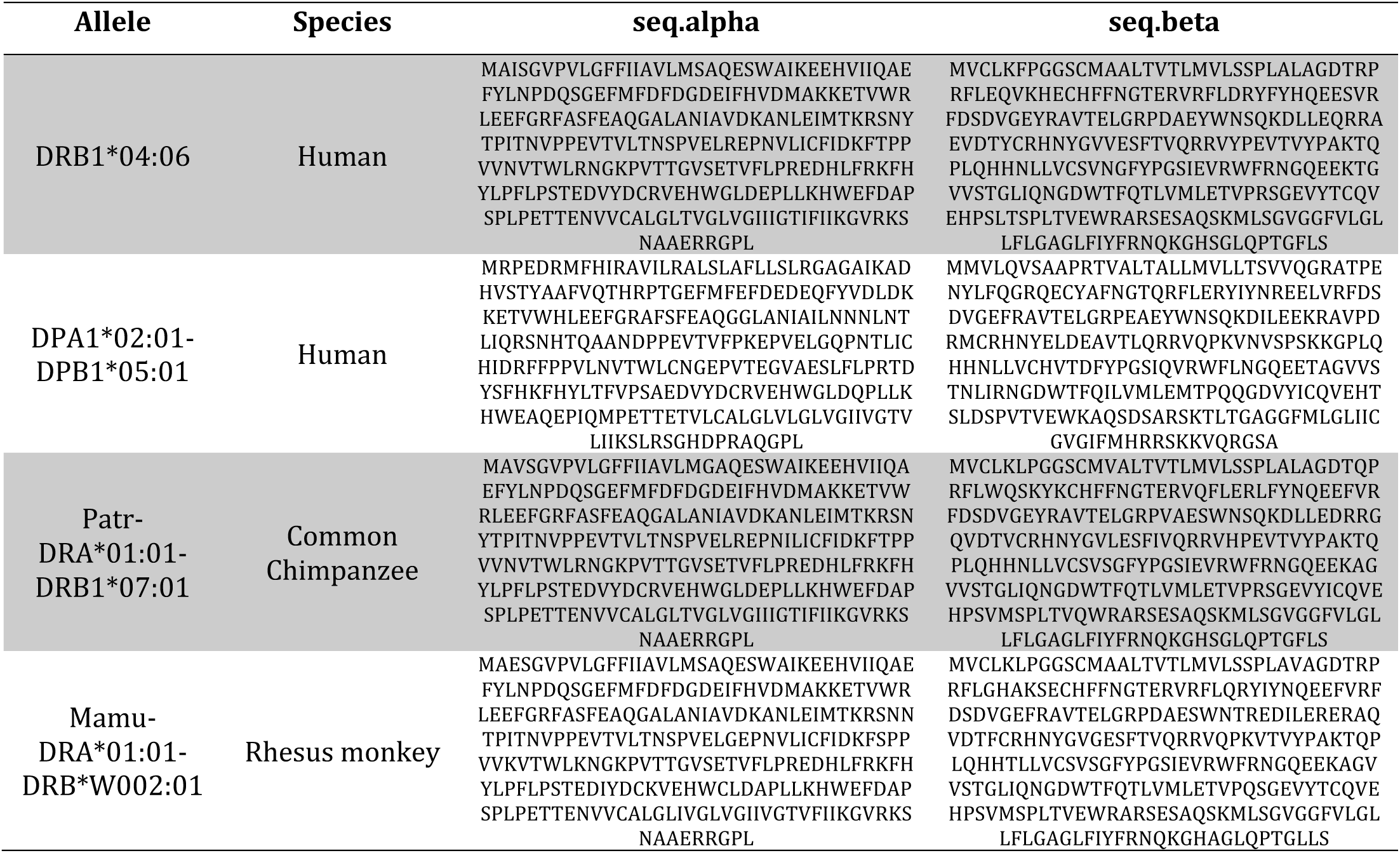
Examples of MHC-II allele sequences that will be analyzed here. The alpha (seq.alpha) and beta (seq.beta) chain sequences of various MHC-II alleles are given, indicating also their species of origin. The full list of alleles used in this book chapter is given in Table S1.

The procedure to follow is the same to predict the motifs and ligands of new MHC-II alleles that have been sequenced in some species or some individuals.

## 3. Methods

### 3.1. Predicting MHC-II binding specificities from MHC-II amino acid sequences

Based on the amino acid sequence of any MHC-II allele, one can use MixMHC2pred webserver to directly derive the motifs describing the binding specificities of this allele. This is a new feature implemented in our server for this book chapter (see Note 1). The procedure to follow is described below.

1. Go to the server page and click on the “*Predict motifs*” tab. This page can also be reached directly at http://mixmhc2pred.gfellerlab.org/motifs (Figure 2A).
2. The input for these predictions is a set of MHC-II sequences given as a tab-delimited text file (Table S1 for example). This file should contain a header row and at least 3 columns: *Allele* (i.e., the name of the MHC-II allele; see Note 2), *seq.alpha* and *seq.beta* (the alpha and beta chain sequences, respectively; see Note 3). Each row corresponds to a different MHC-II allele. Upload this file to the server (Figure 2A – input (1)), either by clicking on the blue box or by dragging-and-dropping the file into the box. See also Note 4 for an alternative input, based on allele names.
3. Click on the button “*Predict allele specificities*” to launch the predictions. Wait a little while.
4. Once the predictions are finished, a new page will open with the results (Figure 2B). On top of this page, you can find a “*Motifs run ID*” that can be used later (Figure 2B – (4); see Note 5 and Section 3.2.1). Below it, there are links to download either the full results including an html page of these or only the PWM definition files (Figure 2B – (5); see Note 6). A table summarizing the user-inputted alleles follows (Figure 2B – (6)) and one can click on an allele name to jump to the part showing the predicted binding specificity motif(s) of this allele (Figure 2B – (7)). When an allele is predicted to possess multiple specificities, the motif from each specificity is returned (as e.g., for DPA1*02:01/DPB1*05:01 in Figure 2B), indicating the binding orientation of ligands towards the given specificity and the predicted fraction of ligands following each specificity.
5. As a quality control check, we advise users to verify the “binding site similarity” (“*BS sim.*”) indicated in the summary table for each allele (Figure 2B - (6)). We usually expect good prediction accuracy for values of this similarity above 0.8 (see section 5 for details). We further advise users to verify the quality of the alignment from their MHC-II sequences in the binding site region. This alignment is present in the “*allelesTable.txt*” file available with the results (obtained when downloading the full results or with the link “*Alleles’ information table*” in Figure 2B - (5)). The presence of many gaps or unusual amino acids with respect to reference sequences likely indicate issues with the inputted MHC-II sequences and can result in lower quality of MixMHC2pred predictions (see also Note 3 and section 5).

Figure 3 shows the binding specificities that have been predicted in this way for some alleles from Table S1. For the human alleles, recent MHC-II peptidomics data was available (***18***). We could therefore compare the predicted motifs with the ones obtained directly from experiments (Figure 3A; see Notes 7 and 8), indicating accurate predictions.

**Figure 3:**
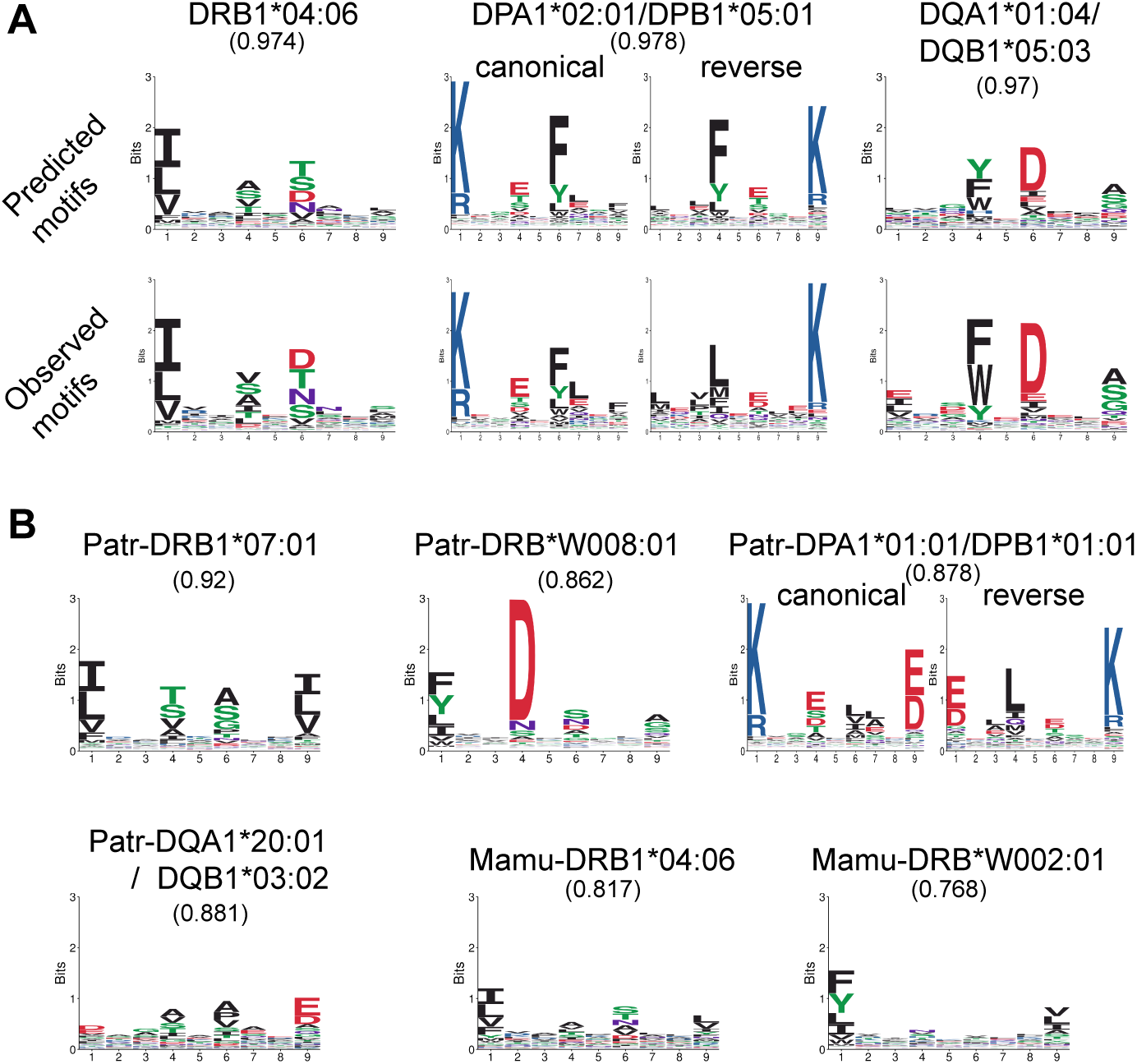
Binding specificities predicted by MixMHC2pred. (A) Comparison between predicted motifs (top row) and those observed in MHC-II peptidomics data from human alleles (bottom row). (B) Motifs predicted for some non-human alleles, including one case with ligands predicted to bind in two orientations. Numbers in parenthesis below the allele names indicate the binding site similarity of the allele.

### 3.2. Predictions of ligands and epitopes

Having determined the binding specificities of MHC-II alleles based on their amino acid sequences, we can now process to predict MHC-II ligands and CD4^+^ T-cell epitopes for these alleles (see Note 9). As an example, we will predict here the presentation of a small set of CD4^+^ T-cell epitopes from common chimpanzee ((***22, 23***); Table 2, see Note 10). We will do predictions for these epitopes with each chimpanzee allele present in Table S1 (alleles starting with “*Patr-*“), as a way of predicting potential ligands from these alleles as well as determining the candidate restriction alleles for these epitopes. Section 3.2.1 explains how to do it based on MixMHC2pred server and section 3.2.2 describes this for the standalone executable version. Finally, section 3.2.3 will compare the predictions to experimental results for this dataset as well as for another study from a different species.

**Table 2:** List of CD4^+^ T-cell epitopes for common chimpanzee alleles, whose restriction allele could be experimentally determined, derived from Woollard et al. (*22*) and Shoukry et al. (*23*). Columns indicate the sequence of the epitope used in our predictions, the minimal epitope sequence and the restricting allele, which were both determined experimentally. See Note 10.

| Peptide | Minimal epitope | Restricting allele |
| --- | --- | --- |
| AIASLMAFTAAVTSP | LMAFTAAVTS | Patr-DRB1*07:01 |
| YTFALWRVSAEEY | ALWRVSAEEY | Patr-DRB1*07:01 |
| YAAQGYKVLVLNPSV | GYKVLVLNPSV | Patr-DRB1*10:01 |
| AATAFVGAGLAGAAI | AFVGAGLAGA | Patr-DRB1*10:01 |
| PFYGKAIPLEVIKG | YGKAIPLEVI | Patr-DRB5*03:10 |

#### 3.2.1 Web application

1. Go to MixMHC2pred server main page (http://mixmhc2pred.gfellerlab.org; Figure 4A).
2. Copy-paste the list of peptides for which you want predictions into the corresponding input box (Figure 4A – (1); see Notes 11 and 12).
3. For this example, the “*Include context encoding*” box should not be ticked (Figure 4A – (2); see Note 11).
4. Give a name for the results file in the corresponding field (Figure 4A – (3)).
5. Enter the list of alleles to consider in the next field (Figure 4A – (4)), e.g., “*Patr-DRA*01:01/DRB1*07:01, Patr-DRA*01:01/DRB1*10:01, Patr-DRA*01:01/DRB5*03:10, Patr-DRA*01:01/DRB*W008:01, Patr-DPA1*01:01/DPB1*01:01, Patr-DQA1*20:01/DQB1*03:02*”.
6. When doing predictions based on user-defined alleles, enter next the Motifs run ID obtained previously (Figure 4A - (5); see Note 5). This ID is automatically populated if the steps from section 3.1 had been done right before.
7. Click on “*Run*”. Predictions will quickly be finished, and the text file of the results can directly be downloaded (Figure 4B; see Note 13).

#### 3.2.2 Executable

1. To run MixMHC2pred executable based on user-defined alleles, first download the PWM definition files that you obtained previously (see section 3.1, step 4; Figure 2B – (5)). This can either be only the PWM definition files or the full results.

a. Uncompress the given .zip file of the results into a folder of your choice.
b. Note the path of the corresponding “*PWMdef*” folder found in these results (we assume below that it is “*path_to_PWMdef_folder*”).
2. Create a text file containing only the peptide sequences to test, one peptide per line (if considering the context, this file should have a 2^nd^ column with the context sequence). This file should not contain any header row. For the example based on Table 2, this file would thus have 5 rows containing: *AIASLMAFTAAVTSP, YTFALWRVSAEEY, YAAQGYKVLVLNPSV, AATAFVGAGLAGAAI, PFYGKAIPLEVIKG*.
3. Open a command window (e.g., a “Terminal” window on Mac OS or a “Command Prompt” on Windows).
4. Run MixMHC2pred: ~~~
path_to_MixMHC2pred/MixMHC2pred.exe -i
  path_to_input_pep/input.txt -o path_to_output/output.txt –
  no_context -f path_to_PWMdef_folder -a <list_of_alleles>
~~~ Where the various “*path_to_…*” are replaced by the corresponding paths, *MixMHC2pred.exe* is used on Windows (*MixMHC2pred* on Mac *OS* and *MixMHC2pred_unix* on Unix), “*input.txt*” gives the name of the input peptide sequence file, “*output.txt*” gives the name of the file where to save the results, and “*<list_of_alleles>*” is replaced by the list of alleles for which we want the predictions, with a space between each allele name (in this example: *Patr_DRA_01_01 DRB1_07_01 Patr_DRA_01_01 DRB1_10_01 Patr_DRA_01_01 DRB5_03_10 Patr_DRA_01_01 DRB_W008_01 Patr_DPA1_01_01 DPB1_01_01 Patr_DQA1_20_01 DQB1_03_02*). Note that for the allele names, the “*”, “:” and “-” should be replaced by “_” and the “/” should be replaced by two “_” (i.e., “ ”): this is to avoid issues with these symbols that have special meanings in command windows. The “*-f path_to_…*” is not needed if predicting for human alleles already available with MixMHC2pred (see Note 9). See Note 14 for further details about the command used above.
5. The results are available at “*path_to_output/output.txt*” for the example command (Figure 4B; see Note 13).

**Figure 4:**
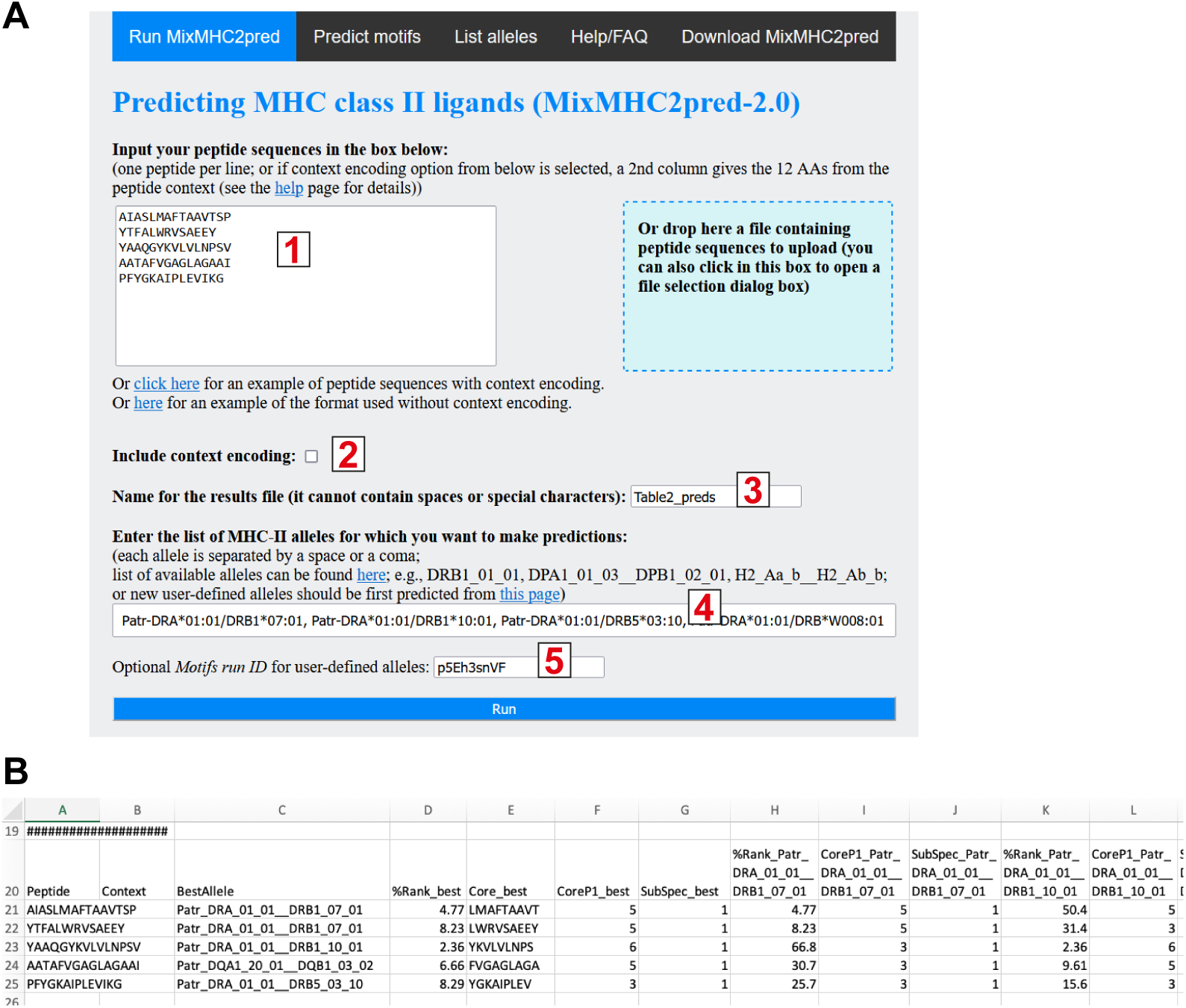
Predicting ligands and epitopes with MixMHC2pred. (A) Main input form to predict peptide presentation for a set of alleles. (B) Snapshot of the result file for predictions with MixMHC2pred.

#### 3.2.3 Analysis of the results

As a first example we compare the results for the small set of chimpanzee CD4^+^ T-cell epitopes used in the step-by-step procedures of sections 3.2.1 and 3.2.2 (Table 2). Results from MixMHC2pred for these peptides with the set of chimpanzee MHC-II alleles tested here are shown in Figure 4B and Figure 5A (see Note 15). These reveal that the positive peptides were well predicted towards their cognate MHC-II allele. Looking at the predicted binding cores of these peptides (Figure 4B), we see that they were always fully included in the minimal epitope determined experimentally in (***22, 23***) (see Table 2), showing that MixMHC2pred correctly identified the binding cores. Comparing the predictions per allele, we note that the experimentally determined epitopes from an allele had clearly better scores than the other peptides (Figure 5A), demonstrating MixMHC2pred’s ability to predict the positives among a set of peptides. Alternatively, analysing the results per peptide, we find that such predictions can also help in determining the restriction allele from the peptides: each peptide had its best score with its experimentally determined restriction allele, except for AATAFVGAGLAGAAI that scored best with Patr-DQA1*20:01/DQB1*03:02 (Figures 4B and 5A). This peptide nevertheless also scored well with Patr-DRB1*10:01, and we cannot exclude that both alleles may bind this peptide and that other CD4^+^ T cells would be able to recognize this peptide when presented by Patr-DQA1*20:01/DQB1*03:02.

**Figure 5:**
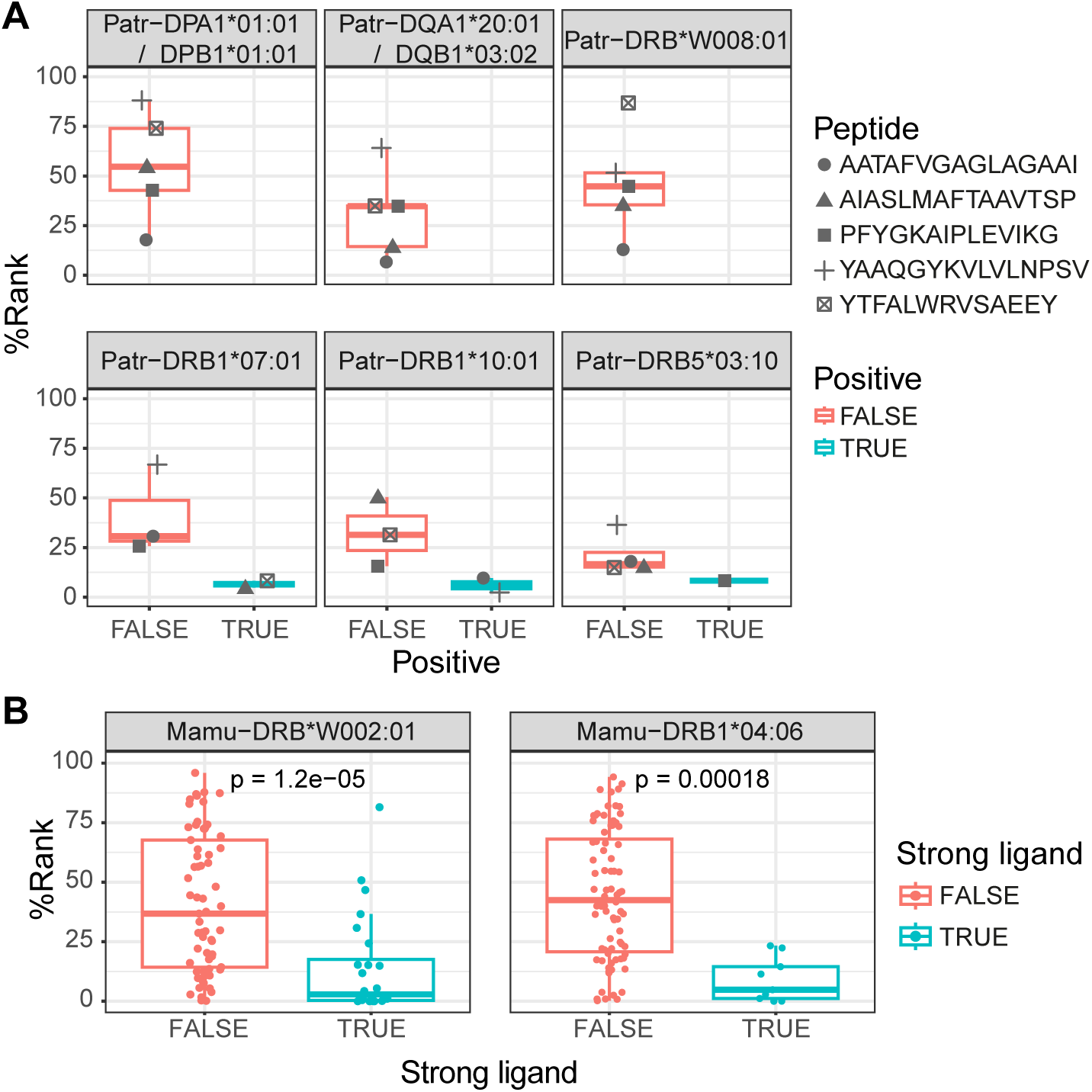
Comparison of predictions to experimental measurements. **(A)** Predicted %Rank scores for various peptides with different chimpanzee MHC-II alleles, compared to their CD4^+^ T-cell immunogenicity determined in (***22*, *23***). **(B)** Predictions for peptides towards rhesus monkey MHC-II alleles from (***24***), comparing the ligands measured as strong binders (K_D_ < 100 nM) for each allele with the rest of peptides. P-values from a two-sided Wilcoxon signed rank test are indicated.

As another example of predictions for MHC-II alleles from a species without any training data, we used a set of peptides tested for binding with rhesus monkey alleles ((***24***); Table S3). Doing predictions for these peptides with MixMHC2pred following the procedure from section 3.2, we observe that the strong binders had significantly better scores than the other peptides (Figure 5B), indicating that MixMHC2pred is able to predict ligands from these rhesus monkey MHC-II alleles as well.

## 4. Notes

1. As described in our publication (***6***), MixMHC2pred is composed of two blocks of neural networks. The first block takes as input the amino acid sequence of an MHC-II allele and predicts its binding specificity (i.e., PWM). The second block takes this PWM as input, as well as the peptide sequences and predicts if the peptides are presented by the corresponding allele. The executable version of MixMHC2pred and main webserver page perform the second block of predictions, based on pre-computed binding specificities (these predictions are described in section 3.2 of this book chapter). In this section 3.1, we describe how to use the webserver to perform the predictions of the first block, to obtain binding specificities for any MHC-II allele. These predictions need to be performed only once and can then be used in the second neural network block.
2. In Table 1, and Table S1, we used the official allele names, but any name would work, as long as it contains only letters, numbers and the symbols “_”, “-“, “:”, “*” and “/”. We adopted the standard practice of not indicating the HLA-DRA allele name, as this gene is not polymorphic contrary to the other HLA genes.
3. For MHC-II alpha and beta chain sequences, the full-length amino acid sequences should ideally be given. Partial sequences should work if the full peptide binding site sequences are present. We also emphasize that the quality of the MHC-II sequence reconstruction can have an impact on the predictions of motifs. As the MHC locus is complex, it may be difficult to precisely determine these sequences in new species, due to low coverage for example. This can potentially render predictions of MHC-II motifs and ligands meaningless. For this reason, we advise to verify the resulting sequence alignment, as described in step 5 from section 3.1.
4. Instead of inputting these allele sequences, if one wants to predict binding specificities from alleles already available in MixMHC2pred, it is possible to list these MHC-II allele names in the corresponding text box, with allele names separated by a space or coma (Figure 2A – input (2)). The list of alleles available can be found at http://mixmhc2pred.gfellerlab.org/alleles.
5. Results from the steps from section 3.1 are stored on the server for about one month and they can be obtained back during this period using the Motifs run ID: either on the “Predict motifs” page to display back the full report (entering this ID in the corresponding text box: Figure 2A – input (3)), or on the main page to predict ligands and epitopes with MixMHC2pred based on user-defined MHC-II alleles (see Section 3.2.1).
6. To predict ligands and epitopes for these user-inputted alleles with the executable version of MixMHC2pred, we need to download these PWM definition files as also explained in section 3.2.2.
7. Note that this MHC-II peptidomics data was absent from the data used to train the current version 2.0.2 of MixMHC2pred and therefore no data corresponding to these 3 *new* human alleles were available for the step to predict motifs with MixMHC2pred.
8. We can use MoDec (***12***) to determine the binding specificities of the alleles present in MHC-II peptidomics data (here the samples were mono-allelic samples, but MoDec works as well on multi-allelic samples). To reproduce these results, we have filtered the peptides found in Stražar et al. (***18***) for the alleles DRB1*04:06, DPA1*02:01/DPB1*05:01 and DQA1*01:04/DQB1*05:03 and copied these into Table S2A-C respectively. We can then determine the binding specificities with the command: ~~~
MoDec.exe -i path_to_pepData/TableS2A.txt -o
  path_to_out_folder/TableS2A --Kmax 3 --MHC2 –makeReport
~~~ Where “*path_to_pepData*” and “*path_to_out_folder*” are replaced by the path where the Table S2A.txt is and the folder where to save the results, respectively. TableS2A can be replaced by TableS2B or TableS2C for the other alleles. For Tables S2A-B, results can take long (around an hour) as these tables correspond to big lists with more than 10 000 peptides each (one could reduce the timing with the option “--nruns 2” for example, which will test fewer initial conditions, possibly missing the optimal motifs but often resulting in similar results, especially in such monoallelic samples). Looking at the report “*path_to_out_folder/TableS2A/TableS2A_report.html*” we see for example that 1 motif was sufficient to describe the binding specificity of the peptides from this allele (DRB1*04:06 for Table S2A). The peptides assigned to other motifs appearing when asking for 2 or 3 motifs correspond to contaminants peptides not showing any clear anchor residue specificity.
9. For this step-by-step example, we started from MHC-II sequences, and we obtained their binding specificities through the steps from section 3.1. But many MHC-II alleles, including non-human alleles (corresponding to more than 80 000 MHC-II heterodimers), are readily available for MixMHC2pred without the need to follow the steps from section 3.1. The list of available alleles is present at the page http://mixmhc2pred.gfellerlab.org/alleles. For these alleles, it is sufficient to use their names when doing the predictions through MixMHC2pred server. If using the executable version, only the human alleles are directly included; for the other species one can download the “.zip” files from above’s page and follow the same procedure than described in section 3.2.2 as for user-defined alleles.
10. The sequences from Table 2 have all been tested experimentally in the studies (***22, 23***) and all were recognized by CD4^+^ T cells when presented by the indicated MHC-II allele. Various additional overlapping sequences have been tested in these studies and we kept here a single sequence per epitope. We do not perform predictions directly against the minimal epitope sequence as MixMHC2pred only returns predictions for peptides longer than 12 amino acids.
11. In this example, we do not consider the peptide context, i.e., we only give the peptide sequences as input, without their context, and we do not tick the option to “*Include context encoding*”. This is because the peptides considered here have been tested as is for binding affinity or immunogenicity. I.e., these peptides have been synthesized and tested directly. The antigen processing starting from full-length proteins and including their cleavage into these peptides was thus not performed by the cell. The context encoding is on the other hand recommended when the goal is to determine the best candidate epitopes starting from a list of overlapping peptides covering some proteins or mutations.
12. Instead of writing the list of peptides to test in the text area box, user can also upload a txt file containing these. In such a case, the file is a simple tab-delimited text file, with a single column if not considering context encoding, or with two columns (1^st^ column: the peptide sequence; 2^nd^ column: the context of the peptide). This file should only contain the peptides, without any header row.
13. The result file is a tab-delimited text file which contains header rows followed by the prediction results: Column (A) indicates the peptide sequence; column (B) is the peptide context (empty in our example as context was not considered); (C) indicates which was the best allele for the given peptide among the set of alleles tested; (D) gives the %Rank of the peptide against this best allele (lower values are better); (E-F) give the predicted binding core against the best allele and starting position of this core sequence; (G) indicates the binding specificity (e.g., a value of “-1” if the peptide is predicted to bind in the reverse orientation against the given allele); (H and following columns) give the same information than previous columns, separately for each allele.
14. (i) Note that the “*path_to_output*” folder should exist before running MixMHC2pred as it will not create it. (ii) The “*--no_context*” option was used as explained in Note 11 above. (iii) The “*path_to_MixMHC2pred/*” is not needed if MixMHC2pred executable has been set in the user path as written in the README file of this tool. (iv) Concerning the *“-f path_to_PWMdef_folder*” option, this tells where to search for the allele definition files of the user-defined alleles (if any). It is possible to include multiple paths separated by a space (and it is possible to copy/paste or move the “*.txt*” files present in one *PWMdef* folder to another one to group these together (if the allele names are unique)). (v) The list of alleles can combine some alleles found in the “*path_to_PWMdef_folder*” with pre-defined alleles from MixMHC2pred (found in the *PWMdef* folder from MixMHC2pred executable). When a same allele name appears in multiple PWMdef folders, the first folder containing it is used (user-defined folders given with the “*-f*” option are searched first).
15. To import the results from MixMHC2pred and make the figures, we used the free software for statistical computing and graphics: R. We used R version 4.2, which can be obtained from https://www.r-project.org and we also used the following R-packages that needed first to be installed within R: *ggplot2* (v3.4.2), *dplyr* (v1.3.3) and *ggpubr* (v0.4.0). Other software could also be used for these analyses.

## 5. Discussion

In section 3.1, we described how the motifs describing ligands of MHC-II alleles can be obtained with MixMHC2pred server. This procedure is straightforward, requiring only the MHC-II allele amino acid sequences as input and returning predictions in usually less than a minute for multiple alleles simultaneously. To our knowledge, MixMHC2pred is the only tool that can directly predict MHC-II motifs based on their amino acid sequences. An additional asset of MixMHC2pred is its ability to accurately predict multiple motifs for alleles accommodating ligands binding in both the canonical and the reverse orientation. An alternative approach to infer motifs is to use the pan-allele predictors, perform predictions for a large set of random peptides (at least 100 000 peptides), select the best scoring peptides (typically the top 1% best scoring peptides), and build the motif based on the predicted binding cores of these peptides. Such an approach is feasible with MixMHC2pred and NetMHCIIpan (***21***) in any species. Other MHC-II ligand predictors accommodating alleles without experimental ligands do not output the predicted binding cores and can moreover only be used for HLA-DR (***25***) or human/mouse (***26***) alleles. This approach is much more laborious, depends on ad hoc thresholds (e.g., top 1%) and predictions are influenced by the origin of the random peptides (e.g., human proteome versus random amino acid choices). We therefore anticipate that the framework adopted in MixMHC2pred and the new interface could considerably facilitate predictions of binding motifs for new MHC-II alleles.

We showed in Figure 3A that MixMHC2pred was able to accurately predict motifs for new human MHC-II alleles (i.e., alleles for which no ligands were available at the time when MixMHC2pred was trained). No additional MHC-II peptidomics data was available to validate similarly our predicted motifs for new non-human MHC-II alleles. Instead, we directly predicted specific ligands and epitopes from various non-human MHC-II alleles, showing high agreement with experimental measurements (Figure 5). We had also previously performed a detailed benchmark of prediction accuracy, including predictions for new species (see Figures 4, 5 and S4C-D in Racle et al. (***6***)). Our results demonstrated that binding specificities for new alleles can generally be well predicted. We have also shown that the prediction accuracy is strongly correlated with the similarity of the binding site of the allele used as input to alleles with known ligands (Figure S4C in (***6***)). This binding site similarity ranges between 0 and 1, with 1 indicating that the allele provided in input has exactly the same binding site as one allele in the training set of MixMHC2pred. The binding site similarity value is indicated for each allele when predicting its motif (Figure 2B – (6)). As a rule of thumb, one can estimate that binding specificities can be faithfully predicted for MHC-II alleles with a binding site similarity above 0.8 to at least one allele with known ligands. Many alleles from species close to human, like monkeys, will fall into this range of binding site similarity values. The same is true for multiple cattle MHC-II alleles owing to the availability of MHC-II peptidomics data from multiple cattle alleles (***14***). Lower accuracy in binding motif and ligand predictions is expected for more evolutionary distant species like amphibians, fishes or reptiles.

Beyond epitope predictions, our analyses showed that MixMHC2pred can be used to help inferring both the minimal epitope sequence and the allelic restriction of known epitopes. Regarding the allelic restriction, predictions for the multiple MHC-II alleles present in a sample can help determining the restriction, as peptides will often score well for only one or few of these alleles, indicating that the given peptide can likely only be restricted by this subset of alleles. Inferring minimal epitope and allelic restriction in this way can be a nice complement to experiments, as these are laborious, requiring testing the epitope recognition when the peptide is presented by different antigen presenting cells sharing some MHC-II alleles, and such cells are not always available, especially for rarer alleles.

One assumption of our predictions is that the overall structure of the binding site is conserved across species, and that the amino acids at the different positions of the binding site can be determined based on sequence alignment. This assumption is supported by the fact that most MHC-II alleles align well with those with known ligands, and by dozens of crystal structures (mainly for human and mouse alleles). However, we cannot exclude that some species may have evolved different binding site architectures. In these cases, our predictions are likely to be of low quality. We therefore encourage users to always verify the quality of the alignment of the MHC-II allele sequences used in input with those of known MHC-II alleles in human (see section 3.1 and Note 3). This is especially important in cases where MHC-II sequences may be of lower quality (lower sequence coverage of the MHC-II locus, poor annotation in less studied species, etc.), since incorrectly reconstructed amino acid sequences will likely lead to spurious predictions.

Altogether, our results showcase the applicability of MixMHC2pred towards predictions of MHC-II binding motifs as well as MHC-II ligands and epitopes for any MHC-II allele based on its amino acid sequence. Considering its accuracy, speed, and ease of use, MixMHC2pred provides a reliable interface to perform such predictions.

## Acknowledgments

DG and JR acknowledge support from the Swiss Cancer Research Foundation (KFS-4961-02-2020).

